# Discovery of bactericides as an acute mitochondrial membrane damage inducer

**DOI:** 10.1101/2021.04.09.439218

**Authors:** Ryan Houston, Yusuke Sekine, Mads B Larsen, Kei Murakami, Derek P Narendra, Bill B Chen, Shiori Sekine

**Author notes:** Corresponding author Shiori Sekine, Mailing address: Bridgeside Point 1, Room 568, 100 Technology Drive, Pittsburgh, PA 15219.

## Abstract

Mitochondria evolved from endosymbiotic bacteria to become essential organelles of eukaryotic cells. The defined lipid composition and structure of mitochondrial membranes are critical for the proper functioning of mitochondria. However, mitochondrial stress responses that help maintain the integrity of mitochondrial membranes against internal or external insults are not well understood. One reason for this lack of insight is the absence of efficient tools to specifically damage mitochondrial membranes. Here, through a compound screen originally aimed at identifying inhibitors of the inner mitochondrial membrane (IMM)-resident protease OMA1, we found that two bis-biguanide compounds, Chlorhexidine and Alexidine, modified OMA1 activity by altering the integrity of the IMM. Interestingly, these compounds are well-known bactericides whose mechanism of action has centered on their damage-inducing activity on bacterial membranes. We found Alexidine binds to the IMM likely through the electrostatic interaction driven by the membrane potential as well as an affinity for anionic phospholipids. Electron microscopic analysis revealed that Alexidine severely perturbated the IMM, especially the cristae structure. Along with this, we observed the altered localization of IMM-resident membrane-shaping proteins, including Mic60. Notably, Alexidine evoked a specific transcriptional/proteostasis signature that was not induced by other typical mitochondrial stressors, highlighting the unique property of Alexidine as a novel mitochondrial membrane stressor. Our findings provide a chemical-biological tool that can induce acute and selective perturbation of the IMM integrity, which should enable the delineation of mitochondrial stress-signaling pathways required to maintain the mitochondrial membrane homeostasis.

## Introduction

Mitochondria are essential and multi-functional organelles of the cell that are involved in energy production, metabolic processes and cellular signaling. Evolutionally, mitochondria evolved from α-proteobacteria that invaded into host eukaryotic cells. Mitochondria are surrounded by two membranes; the outer mitochondrial membrane (OMM) and the inner mitochondrial membrane (IMM). Both membranes have a characteristic phospholipid composition and structure. Reflecting the endosymbiotic origin of mitochondria, the IMM shares similarity with the lipid composition of bacterial membranes including high levels of cardiolipin (CL) [1–3]. CL is a non-bilayer forming phospholipid that destabilizes the lipid order in bilayers and induces high membrane curvature. Together with phosphatidylethanolamine (PE), which has similar biophysical properties as CL, these non-bilayer lipids make up approximately ∼50% of the phospholipids in the IMM and are more abundant in this membrane than in any other cellular membranes [1–3]. CL and PE contribute to the characteristic highly-folded structure of the IMM as represented by cristae, the concave membrane structure of the IMM [4]. Among the biological membranes in the cells, the IMM is known to possess the highest protein density, allowing various essential bioenergetic reactions to occur. The activities of the IMM-embedded enzymes, including the OXPHOS proteins, rely on the defined lipid composition and structures of the IMM [4]. Therefore, the integrity of the IMM must be carefully monitored and maintained in the face of internal or external insults.

Various mitochondrial stress responses that maintain healthy mitochondrial network have been discovered [5]. These include mitophagy, an autophagic degradation of damaged mitochondria [6], as well as the mitochondrial unfolded protein response (mtUPR) [7], that up-regulates a specific transcriptional program to relieve mitochondrial proteotoxic stress. Importantly, the existence of small molecules that can mimic a distinct type of mitochondrial damage has significantly contributed to the discovery and understanding of these crucial mitochondrial stress-signaling pathways. For example, the mitochondrial protonophore carbonyl cyanide m-chlorophenyl hydrazone (CCCP) and the combination of OXPHOS inhibitors, Antimycin and Oligomycin, have been widely used for the mechanistic analysis of PINK1/Parkin-mediated mitophagy [8–10]. The treatment of the cells with these compounds results in mitochondrial membrane potential loss, one of the hallmarks of OXPHOS dysfunction, triggering mitophagy. In mammalian cells, the mtUPR can be induced by CDDO, an inhibitor of the matrix-resident protease LONP, or GTPP1, a mitochondrial HSP90 inhibitor [11]. Moreover, mitochondrial proteotoxic stress can be induced by Actinonin [12, 13], an inducer of mito-ribosome stalling that results in the blockade of mitochondrial protein translation. A series of mitochondrial import blockers, so-called MitoBloCKs, can impair mitochondrial protein import pathways [14, 15]. Recent omics analyses of the mammalian cells that were treated with these mitochondrial stressors revealed that many of these stressors commonly activate the integrated stress response (ISR) [16]. The ISR induces the expression of particular cytoprotective genes through the activation of the transcription factor (ATF4), suggesting the existence of an intimate mitochondria-nuclear communication to activate the proper stress response following specific mitochondrial stressors. Recent studies using cell genetic screens for genes involved in the mitochondrial stressor-induced ISR revealed that mitochondrial proteolysis plays a critical role in activating the ISR [17, 18]. These examples clearly highlight the importance of chemical compounds that can induce a specific mitochondrial stress in identifying and analyzing mitochondrial stress-signaling pathways.

Recent identification of sets of lipid synthesis enzymes and lipid transfer proteins significantly advanced our understanding of lipid metabolism within mitochondria [2, 3]. However, it is still not known how mitochondrial membrane homeostasis is preserved under conditions that disturb mitochondrial membrane integrity. This is partly due to a lack of established compounds that can specifically perturb the phospholipid environment of mitochondrial membranes.

Here, through our unbiased small-compound screen that targeted the IMM-integral protease OMA1, we found that two small-compounds, Chlorhexidine and Alexidine, that acutely disrupted the integrity of mitochondrial membranes and thereby secondarily alter OMA1 activity. Interestingly, these compounds are known as bactericides that have damage-inducing activities on bacterial membranes. Our biochemical analyses revealed that Alexidine had an affinity to mitochondrial membranes and particularly damage the cristae membranes in the IMM. Moreover, we found that the Alexidine-treatment induced characteristic transcriptional and proteostatic signatures that were not observed with other typical mitochondrial stressors. Our discovery therefore provides a unique chemical-biological tool that can acutely and selectively perturb membrane homeostasis in the IMM.

## Results

### Compound screen of OMA1 inhibitors identifies bactericides

In heathy mitochondria which maintain mitochondrial membrane potential, PTEN-induced kinase 1 (PINK1) is cleaved by the IMM-resident protease PARL just after the mitochondrial import of PINK1 [19] (Supplementary Fig. 1A, left panel). The cleaved product of PINK1 is retro-translocated into the cytosol for proteasomal degradation [20]. In contrast, mitochondrial depolarization induces the mitochondrial import arrest of PINK1, which results in the accumulation of the full-length form of PINK1 and its kinase activation on the OMM of damaged mitochondria [9, 21, 22] (Supplementary Fig. 1A, middle panel). Activated PINK1 promotes the autophagic elimination of damaged mitochondria, so-called mitophagy, cooperatively working with the cytosolic E3 ligase Parkin [6]. Mutations in PINK1 or Parkin cause recessive early-onset Parkinson’s disease (PD), suggesting a protective role of mitophagy in PD pathogenesis [6]. We have previously reported that several PD-related PINK1 mutants are insensitive to the mitochondrial stress-dependent import arrest and fail to accumulate in the OMM [23, 24]. While these PINK1 mutants are cleaved by PARL in a similar way to PINK1 wild-type (WT) in healthy mitochondria, the mis-sorted PINK1 mutants in depolarized mitochondria are instead cleaved by another IMM protease OMA1 and are subsequently subjected to proteasomal degradation [23] (Supplementary Fig. 1A, right panel). PINK1 (C125G) is one of these import arrest-defective PD-related PINK1 mutants. We stably expressed PINK1 (C125G)-EYFP in PINK1 knockout (KO) cells and confirmed our previous findings by immunoblotting. PINK1 protein was not observed under steady state conditions (Fig. 1A, lane 1 and 4). Treatment with the proteasome inhibitor MG132, however, resulted in the accumulation of the cleaved form of PINK1 (C125G)-EYFP (Fig. 1A, lane 3 and 6), indicating constitutive proteasomal degradation of PINK1 (C125G)-EYFP after PARL-mediated cleavage in healthy mitochondria. Importantly, PINK1 (C125G)-EYFP could not accumulate in response to CCCP in control siRNA-treated cells (Fig. 1A, lane 2), while the full-length form of PINK1 (C125G)-EYFP did accumulate when OMA1 expression was suppressed by an OMA1-specific siRNA (Fig. 1A, lane 5). Thus, in depolarized mitochondria, PINK1 (C125G)-EYFP is degraded through OMA1.

**Figure 1.**
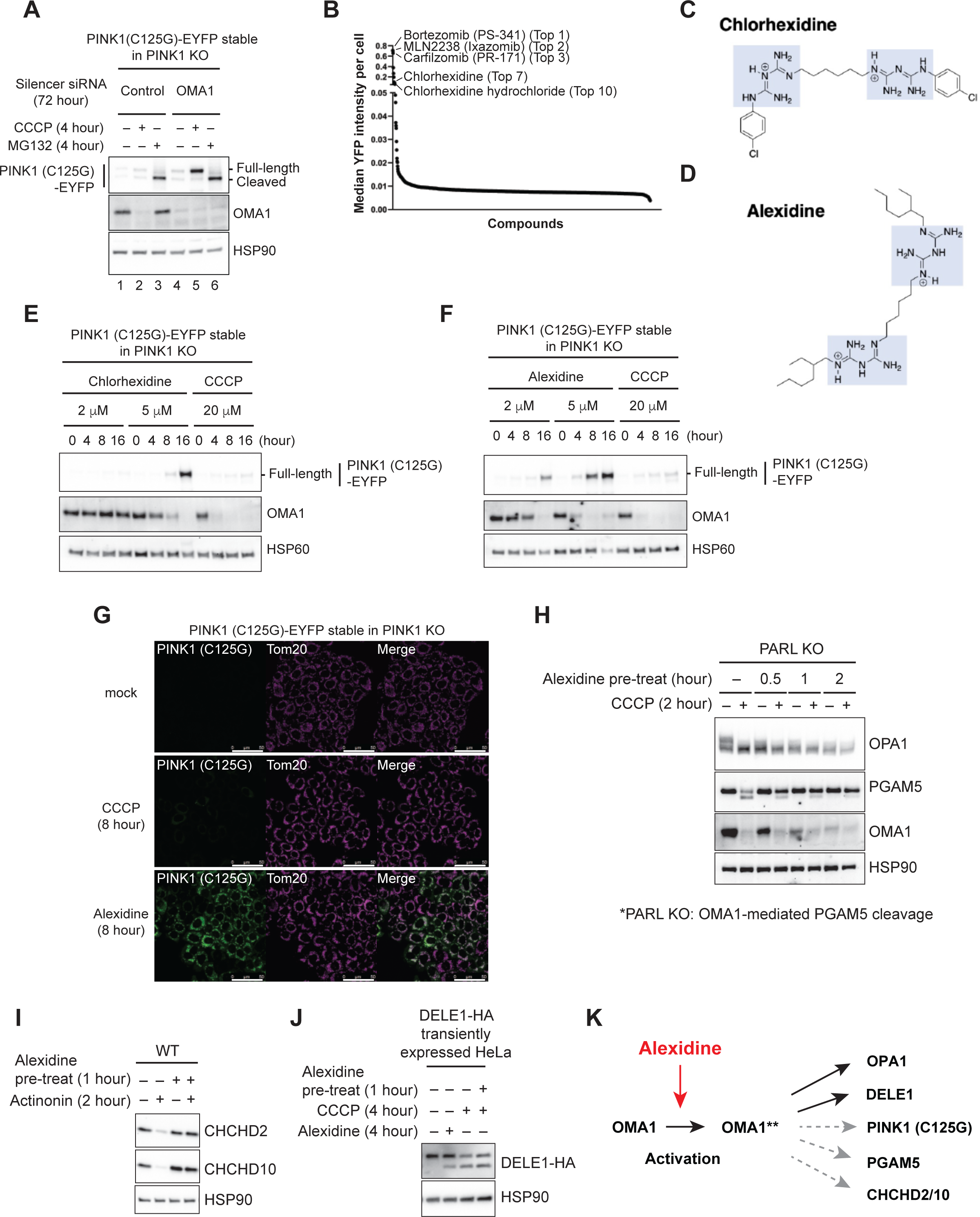
Identification of bactericides as a substrate-dependent OMA1 inhibitor. (A) PINK1 knockout (KO) HeLa cells stably expressed PINK1 (C125G)-EYFP were transfected with control or OMA1 siRNA. After 72 hours, cells were treated with 20 μM CCCP or 10 μM MG132 for 4 hours. The lysate was analyzed by SDS-PAGE. (B) PINK1 KO HeLa cells stably expressed PINK1 (C125G)-EYFP were seeded on 384 well plate, and were treated with each compound (5 μM, FDA-approved compounds). After 18 hours, cells were treated with 20 μM CCCP for 4 hours. EYFP fluorescence intensity of each well was measured by the high content image analyzer. (C, D) The chemical structure of a hit compound, Chlorhexidine (C), and a similar compound, Alexidine (D). Guanidium groups that have delocalized positive charges are highlighted in blue. (E, F) PINK1 KO HeLa cells stably expressed PINK1 (C125G)-EYFP were treated with the indicated drugs for the indicate time period, and the lysate was analyzed by SDS-PAGE. (G) HeLa cells stably expressed PINK1 (C125G)-EYFP were treated with 20 μM CCCP or 5 μM Alexidine for 8 hours, and were subjected to immunocytochemistry (ICC). Tom20 was utilized as a mitochondrial marker. Scale bars; 50 μm. (H) PARL KO HeLa cells were pre-treated with Alexidine for the indicated time period, and after, treated with 20 μM CCCP for 2 hours. Note that in PARL KO cells, the CCCP-dependent PGAM5 cleavage is mediated by OMA1. The lysate was analyzed by SDS-PAGE. (I) WT HeLa cells were pre-treated with 5 μM Alexidine for 1 hour, and after, treated with 150 μM Actinonin for 2 hours. The lysate was analyzed by SDS-PAGE. (J) HeLa cells transiently expressed with DELE1-HA was pre-treated with 5 μM Alexidine for 1 hour, and after, treated with 20 μM CCCP for 4 hours. The lysate was analyzed by SDS-PAGE. (K) Alexidine showed the inhibitory effect on the OMA1-mediated proteolysis in a substrate-dependent manner.

We realized this assay system might allow for the discovery of OMA1 inhibitors since the level of PINK1 (C125G)-EYFP under depolarized conditions was dependent on OMA1 activity. We therefore next measured the intensity of the YFP signal of PINK1 (C125G)-EYFP after CCCP-treatment using a high-content image analyzer. The Z’-factor (a factor that can evaluate the effectiveness of a high-throughput screening) calculated by the YFP signal intensity derived from negative control siRNA vs. OMA1 siRNA yielded a significantly high value (Z’-factor=0.74) (Supplementary Fig. 1B), indicating the potential robustness of our screening system (Z’<0.0, not a suitable assay; 0.0<Z’<0.5, a marginal assay; 0.5<Z’<1.0, an excellent assay; Z’=1.0, ideal assay) [25]. The initial screen was performed with an FDA-approved library consisting of approximately 1,100 compounds (Supplementary table 1). The top 3 hit compounds that induced the accumulation of PINK1 (C125G)-EYFP were proteasome inhibitors (Fig. 1B and Supplementary table 1), confirming the degradation of PINK1 (C125G)-EYFP occurs through proteasome. Among several other hit compounds, we focused on Chlorhexidine and Chlorhexidine hydrochloride, which were independently identified within top 10 hits (Fig. 1B and Supplementary table 1). Chlorhexidine is a bis-biguanide compound (Fig. 1C) that clinically is used as a bactericide, particularly in handwashing and oral care products [26, 27]. There is a similar bis-biguanide compound called Alexidine (Fig. 1D) [26]. These compounds share structural similarities in that they contain symmetrical biguanide units tethered by a long alkyl chain. Strikingly, the treatment with Chlorhexidine or Alexidine, but not CCCP, significantly promoted PINK1 (C125G) accumulation in a dose- and time-dependent manner (Fig. 1E and F). While both Chlorhexidine and Alexidine lowered the mitochondrial membrane potential similar to CCCP (Supplementary Fig. 1C), these observations suggest a membrane depolarization-independent mechanism for the PINK1 (C125G) accumulation by these bactericides. Alexidine was chosen for further study, as it showed a stronger stabilization activity for PINK1 (C125G) than did Chlorhexidine. Alexidine, but not CCCP, induced the accumulation of PINK1 (C125G) on mitochondria (Fig. 1G). These results suggest that the identified bactericides somehow prevented OMA1-mediated PINK1 (C125G) degradation.

### Alexidine demonstrates a substrate-dependent inhibition of OMA1-mediated proteolysis

OMA1 is a stress-responsive protease whose proteolytic activity is enhanced in response to mitochondrial damage including CCCP-induced mitochondrial depolarization [28, 29]. OMA1 activation is achieved by its self-cleavage that eventually leads to the complete degradation of OMA1 [30, 31]. Thus, the degradation of OMA1 indicates its activation. Although Chlorhexidine and Alexidine appeared to inhibit the OMA1-mediated degradation of PINK1 (C125G) (Fig. 1E and F, PINK1 panels), we noticed that the autocatalytic degradation of OMA1 was enhanced in response to these bactericides as observed under CCCP-treated conditions (Fig. 1E and F, OMA1 panels). Our previous report suggested that the OMA1 inhibition not only prevented the degradation of PINK1 (C125G) but also restored its kinase activity and thus the subsequent mitochondrial recruitment of Parkin [23]. However, treatment with Alexidine did not restore the kinase activity of PINK1 (C125G) (Supplementary Fig. 1D, lane 7 in a Phos-tag gel).

These results raised the possibility that the bactericides-mediated stabilization of PINK1 (C125G) cannot simply be attributed to the inhibition of OMA1 proteolytic activity. Therefore, we examined the cleavage of other known OMA1 substrates following Alexidine treatment. We tested the dynamin-like GTPase OPA1 [28, 29], and the mitochondrial protein phosphatase PGAM5 [32, 33]. It is known that both of these mitochondrial proteins are cleaved in response to CCCP. While the CCCP-induced OPA1 cleavage mostly depends on OMA1 [28, 29], the CCCP-induced PGAM5 cleavage redundantly depends on both OMA1 and PARL [32, 33]. To examine the effect of Alexidine on the OMA1-mediated PGAM5 cleavage, we tested whether Alexidine pre-treatment could inhibit CCCP-dependent PGAM5 cleavage in PARL KO cells. We observed that in PARL KO cells, Alexidine pre-treatment inhibited CCCP-induced PGAM5 cleavage in a time- and dose-dependent manner (Fig. 1H and Supplementary Fig. 1E, PGAM5 panels). In contrast, CCCP-dependent OPA1 cleavage was still observed even with Alexidine pre-treatment (Fig. 1H and Supplementary Fig. 1E, OPA1 panels). These results suggest that Alexidine prevented OMA1-mediated PGAM5 cleavage, but not OMA1-mediated OPA1 cleavage. Recently, three other OMA1 substrates were reported; CHCHD2, CHCHD10 and DELE1. CHCHD2 and CHCHD10 are degraded by OMA1 after treatment with Actinonin, a mitochondrial translation inhibitor [34]. DELE1 is cleaved by OMA1 in response to mitochondrial damage including CCCP, which induces the ISR through activation of HRI, one of the eIF2α kinases [17, 18]. Again, Alexidine showed a distinct effect on OMA1-mediated proteolysis depending on the particular substrate. Alexidine pre-treatment inhibited OMA1-mediated degradation of CHCHD2 and CHCHD10 (Fig. 1I), while it failed to inhibit OMA1-mediated cleavage of DELE1 (Fig. 1J). Collectively, these results suggest that Alexidine is not a simple OMA1 inhibitor (rather, it activates OMA1 itself), but it inhibits the OMA1-mediated proteolysis in a substrate-dependent manner (Fig. 1K).

### Alexidine has an affinity for the IMM

We next tried to identify a target of Alexidine to address the underlying molecular mechanism of the observed substrate-specific action of Alexidine on OMA1-mediated proteolysis. In addition to antimicrobial properties, Chlorhexidine and Alexidine are also reported as inhibitors of PTPMT1 [35], a mitochondrial matrix-localized phosphatase that dephosphorylates phosphatidylglycerol-phosphate (PGP), an essential intermediate in cardiolipin (CL) biosynthesis [36, 37]. However, knockdown (KD) of PTPMT1 in our PINK1 (C125G)-EYFP stable HeLa cells did not promote PINK1 (C125G) stabilization (Supplementary Fig. 2, lane 5), suggesting that Alexidine appears to have a different target in this context.

The proposed mechanism of action of Chlorhexidine as a bactericide centers on its bacterial membrane damage-inducing ability through its interaction with phospholipids [26, 27]. Alexidine also has a similar activity on the bacterial membranes [26]. Guanidinium groups of these compounds possess delocalized positive charges at physiological pH [38] (Fig. 1C and D). The delocalized positive charges have higher lipophilicity compared with groups that have localized charges, which is considered to confer the efficient binding ability of Chlorhexidine and Alexidine to phospholipids, together with their long alkyl chain between two symmetric guanidinium groups [26, 27]. These observations led us to examine the effect of Alexidine on the phospholipids in the IMM. We first tested a phospholipid dye 10-N-Nonyl acridine orange (NAO) staining with or without Alexidine-treatment. NAO is a lipophilic and positively charged molecule that is often utilized to stain CL in mitochondria [39, 40], and is also to monitor anionic phospholipids in bacterial membranes [41]. In eukaryotic cells, it is widely known that NAO selectively accumulates in the IMM [39, 40]. We found that treatment with Alexidine but not CCCP dramatically reduced mitochondrial NAO staining (Fig. 2A). NAO staining was rapidly lost after drug-treatment (Fig. 2B). In contrast, the fluorescent intensity of Su9-mCherry (matrix marker) was not affected, indicating that mitochondria themselves were still present (Fig. 2B). These results indicate that Alexidine has effects on the IMM phospholipids. To directly evaluate the binding affinity of Alexidine to phospholipids, we examined the effect of Alexidine on *in vitro* binding between NAO and anionic phospholipid species coated on microplate wells [42, 43]. Pre-incubation with Alexidine was found to reduce the fluorescent intensity derived from NAO bound to anionic phospholipids in a dose-dependent manner (Fig. 2C). Consistent with the NAO staining in cells (Fig. 2A), CCCP did not reduce NAO intensity in this assay (Fig. 2C). These results suggest that Alexidine has an affinity to anionic phospholipids and competes with NAO to bind to these IMM phospholipids.

**Figure 2.**
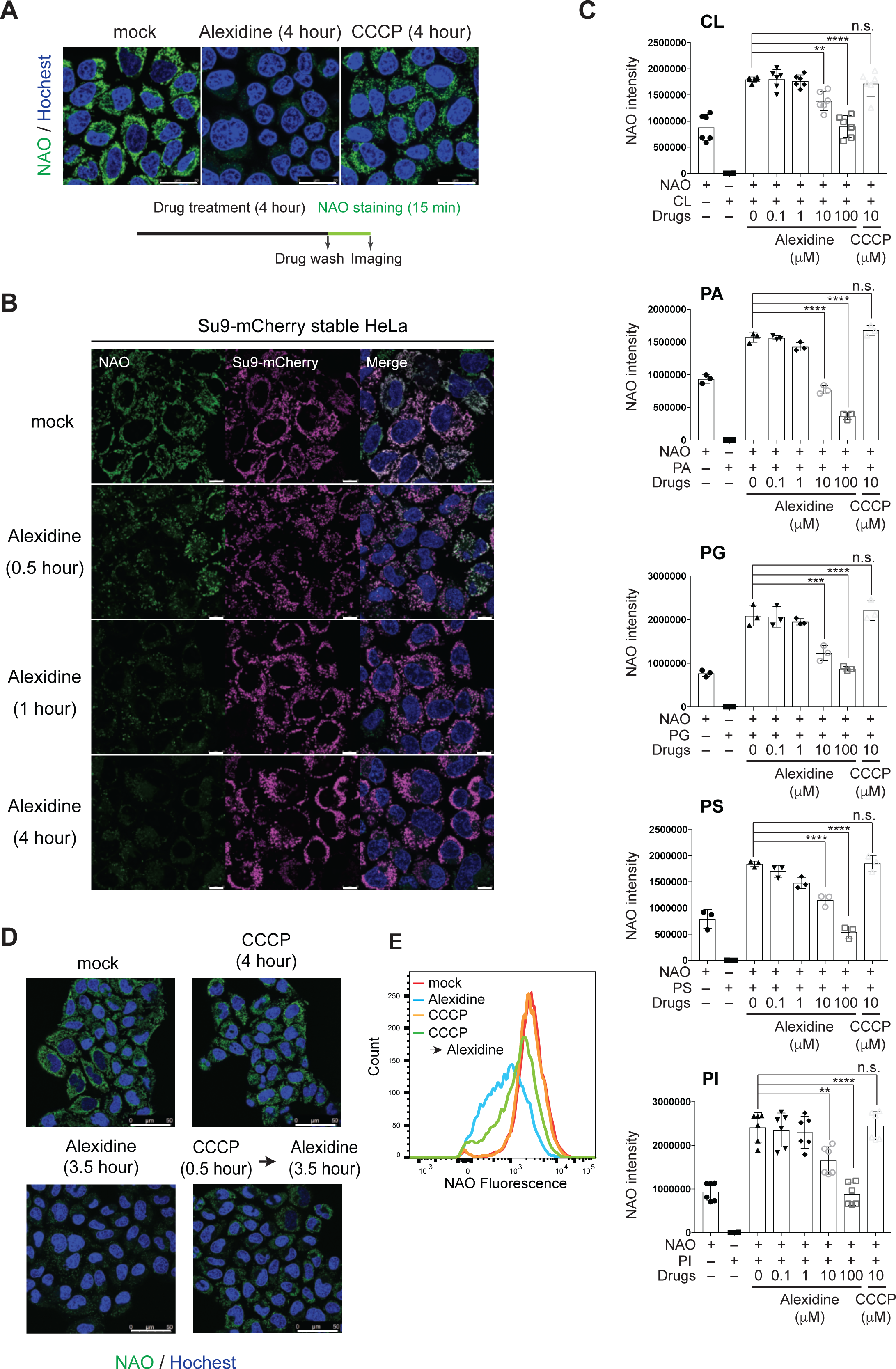
Alexidine has an affinity for the IMM. (A and B) WT HeLa cells or HeLa cells stably expressed Su9-mCherry (a matrix marker) were treated with 5 μM Alexidine or 20 μM CCCP for the indicated time period. After the drug-treatment, cells were washed with phosphate-buffered saline (PBS) for twice, and were stained with NAO. NAO staining was analyzed by live-cell imaging. Scale bars; 25 μm (A) and 10 μm (B). (C) NAO fluorescence intensity (Ex 485nm /Em 535 nm) in individual wells of a microtiter plate which were coated with or without the indicated phospholipid species was measured by microtiter plate reader. Alexidine or CCCP was added at the indicated concentration for 30 min before the NAO staining. Data are shown as mean ± S.D. (n=3 or n=6 per condition). **P<0.01, ***P<0.001, and ****P<0.0001 (One-way ANOVA followed by Turkey’s multiple comparison). (D) WT HeLa cells were treated with 5 μM Alexidine, 20 μM CCCP or the combination of these drugs for the indicated time period. After the drug-treatment, cells were washed with phosphate-buffered saline (PBS) for twice and were stained with NAO. NAO staining was analyzed by live-cell imaging. Scale bars; 50 μm. (E) FACS analysis of NAO fluorescence intensity in (D).

Intriguingly, the aforementioned chemical property of guanidium groups is also often utilized to target drugs to mitochondria, because it is known to preferentially accumulate in the IMM which has an electrochemical gradient [44, 45]. Therefore, we tested whether mitochondrial depolarization prevents the action of Alexidine on the IMM. CCCP-pretreatment partially attenuated the Alexidine-induced reduction of NAO staining (Fig. 2D and E). These results suggest that the mitochondrial membrane potential can be a primary driving force for the mitochondrial targeting of Alexidine, and the high lipophilicity of Alexidine promotes the accumulation of this compound in the hydrophobic lipid environment of the IMM.

### Alexidine induces an acute perturbation of IMM integrity

Since Alexidine appeared to interact with the IMM phospholipids, we investigated the effects of Alexidine on IMM structure and on IMM-shaping proteins. The IMM is structurally subdivided into two domains; the inner boundary membrane (IBM) where the IMM is in close proximity with the OMM, and the cristae, bag-like structures where the IMM invaginates into the matrix [4, 46]. These two IMM domains are connected by narrow, neck-like structures called cristae junctions (CJs) [4, 46] (Fig. 3A). To examine IMM structure, we performed an electron microscopic (EM) analysis of mitochondria with or without Alexidine-treatment. The EM images clearly revealed that the IMM structure was severely disrupted after Alexidine-treatment (Fig. 3A, right panels). The alteration of the cristae membrane was the most striking feature. The cristae membrane was pinched-off from the IBM and often appeared bunched up in an onion-like ball in the matrix. Mitochondria were swollen and the matrix content seemed to be diluted. However, the OMM and the IBM did not exhibit apparent morphological changes and remained in place, suggesting that Alexidine may particularly influence the cristae membranes and CJs. In contrast, CCCP-treatment only displayed a mild disturbance in the IMM structure (Fig. 3A, middle panels).

**Figure 3.**
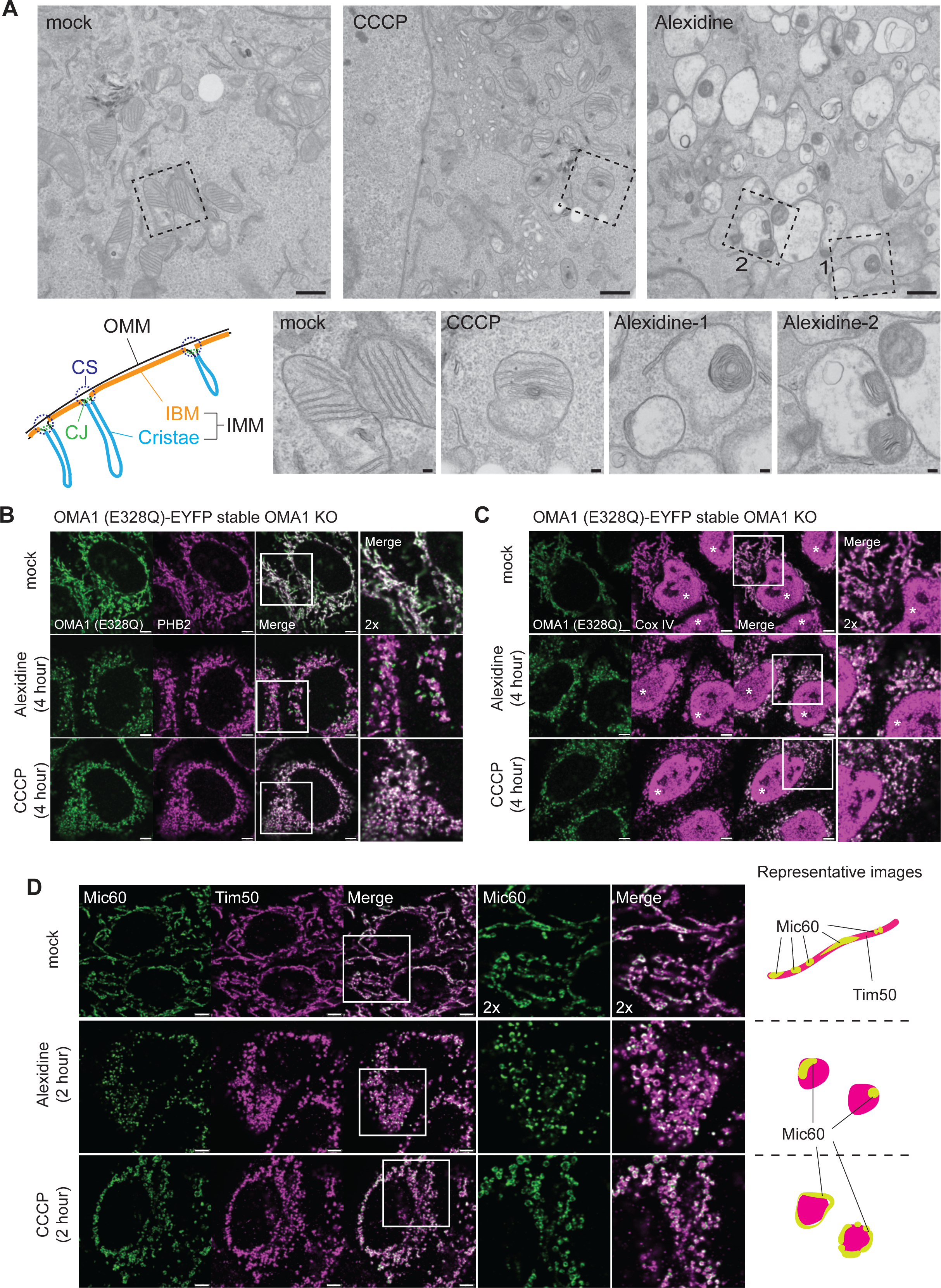
Alexidine induces an acute perturbation of IMM integrity. (A) The electron microscopic (EM) images of the WT HeLa cells that were treated with DMSO, 20 μM CCCP, or 5 μM Alexidine for 4 hours. Scale bars; 800 nm (upper panels), 100 nm (lower panels). (B, C) OMA1 KO cells stably expressed OMA1 (E328Q)-EYFP were treated with 5 μM Alexidine or 20 μM CCCP for 4 hours and were subjected to ICC. Cox IV was used as a typical IMM marker protein (C). *; non-specific signal. Scale bars; 5 μm. (D) WT HeLa cells were treated with 5 μM Alexidine or 20 μM CCCP for 2 hours and were subjected to ICC. Tim50 was used as a typical IMM marker protein. Scale bars; 5 μm.

We next examined the effects of Alexidine on proteins that help shape the IMM. The PHB complex is a heteromeric complex which consists of the closely related IMM-localized proteins PHB1 and PHB2, and is predicted to form a ring-like structure in the IMM [47]. It has been shown that the PHB complex may regulate the CL/PE-enriched microdomains in the IMM [48–50]. At the same time, the PHB complex is also known to regulate several IMM-resident proteases including m-AAA [51], OMA1 [52, 53], and PARL [54]. All of these proteases are single- or multi-spanning membrane proteins, suggesting the importance of the PHB complex-organized lipid micro-environments in the regulation of their proteolytic activities. Because we observed substrate-specific effects of Alexidine on OMA1-mediated proteolysis (Fig. 1K), we next examined the localization of OMA1 and PHB2. To prevent the stress-dependent auto-catalytic degradation of OMA1, we stably expressed an OMA1 protease activity-dead mutant, OMA1 (E328Q) [23], in OMA1 KO cells. The localization of OMA1 and PHB2 mostly overlapped under steady state and CCCP-treated conditions (Fig. 3B, upper and lower panels). However, under Alexidine-treated conditions, the localization of these proteins diverged, with part of the OMA1 pool now segregated from the PHB2-positive IMM (Fig. 3B, middle panels). When the IMM was stained with Cox IV, a subunit of the cytochrome c oxidase complex, the localization of OMA1 overlapped with the Cox IV-positive IMM even under Alexidine-treated conditions (Fig. 3C, middle panels), indicating that OMA1 still exists in the IMM.

The MICOS complex is located at CJs where it stabilizes membrane curvature and forms contact sites (CSs) between the OMM and the IMM [46] (Fig. 3A). It is reported that the MICOS complex genetically interact with the CL synthesis pathway [55, 56], and that some components (Mic60 and Mic27) of the MICOS complex directly bind to CL *in vitro* [57, 58]. Among the 7 components of the metazoan MICOS complex [46], we examined Mic60 localization before and after the Alexidine-treatment. Under the resolution of conventional confocal microscopy, Mic60 shows a uniform distribution along the mitochondrial string at steady state conditions (Fig. 3D, upper panels). However, after Alexidine-treatment, Mic60 was localized in a restricted region of each fragmented mitochondrion and showed an intense, bright, puncta-like localization within the IMM (Fig. 3D, middle panels). In contrast to the Alexidine-treated cells, Mic60 was uniformly distributed in each fragmented mitochondrion after CCCP-treatment (Fig. 3D, lower panels), indicating that Mic60 puncta formation is specifically induced by Alexidine. Taken together, these observations suggest that Alexidine induced acute redistributions of IMM-resident proteins along with the perturbation of IMM subdomain integrity.

### Alexidine evokes a unique transcriptional/proteostasis signature

From the observations above, we hypothesized that Alexidine could be used as an acute mitochondrial membrane damage inducer. Therefore, we decided to characterize the cellular response elicited by the Alexidine-induced mitochondrial membrane perturbation. For this purpose, we performed TMT-based quantitative proteomics (Supplementary table 2). As described so far, Alexidine induced mitochondrial alterations that were distinct from CCCP-treatment. In order to identify the proteins whose expression was specifically changed in response to the Alexidine-induced mitochondrial membrane stress, we compared three different conditions: mock, Alexidine and CCCP-treatment. Gene Ontology analysis for proteins which were significantly changed in the Alexidine-treated cells showed a significant enrichment of mitochondria-related proteins (Fig. 4A), suggesting that Alexidine preferentially affects mitochondria among several other organelles. Many proteins were commonly upregulated or downregulated in both the Alexidine and CCCP-treated cells (Fig. 4B), which may be attributed to the observation that Alexidine also induces mitochondrial depolarization at almost the same level as CCCP (Supplementary Fig. 1C). Notably, the expression of some proteins was specifically altered in the Alexidine-treated cells. Twenty-seven proteins were specifically identified as down-regulated proteins in Alexidine-treated cells (fold change < 0.8, T-TEST q-value < 0.05), (Fig. 4C and Supplementary table 2). Among these, 13 proteins were mitochondrial proteins (Fig. 4D). These include OXPHOS proteins, proteins which are involved in Coenzyme Q biosynthesis, and PTPMT1. As up-regulated proteins (fold change > 1.5, T-TEST q-value < 0.05), only 4 non-mitochondrial proteins were specifically up-regulated in response to Alexidine (Fig. 4E and Supplementary table 2). These include metallothioneins (MTs) and heme oxygenase 1 (HMOX1), an essential enzyme in heme catabolism [59] (Fig. 4F). We next sought to confirm the Alexidine-specific expression changes obtained from the proteomics analysis. Immunoblotting analysis confirmed that HMOX1 was up-regulated while COA7, COX17 and PTPMT1 were down-regulated upon Alexidine-treatment (Fig. 4G). Strikingly, these changes were only observed by Alexidine-treatment, and not by other well-known mitochondrial stressors such as CCCP, Rotenone (a Complex I inhibitor), Actinonin or CDDO (Fig. 4G). We confirmed that each mitochondrial stressor was working in our system by monitoring the OPA1 cleavage and the OMA1 activation on immunoblotting (Fig. 4G), and mitochondrial ROS production by FACS (Supplementary Fig. 3A). CDDO induced the up-regulation of mtHSP60 transcription, one of the targets of mtUPR [11], but Alexidine did not show any enhancement of mtHSP60 transcription (Supplementary Fig. 3B). RT-PCR analysis revealed that the upregulation of HMOX1 and MT2A was induced at the transcriptional level (Fig. 4H), indicating the existence of a mitochondrial-nuclear communication in response to Alexidine-treatment. In contrast, the mRNA level of COA7 or COX17 did not change after Alexidine-treatment, indicating that Alexidine likely induces the post-transcriptional degradation of these proteins (Fig. 4H). LONP knockdown prevented the Alexidine-induced downregulation of PTPMT1 (Supplementary Fig. 3C), suggesting that at least PTPMT1 is degraded within the mitochondria. These results suggest that Alexidine can evoke unique mitochondrial responses that are not induced by other typical mitochondrial stressors, presumably through the perturbation of the IMM integrity (Supplementary Fig. 3D).

**Figure 4.**
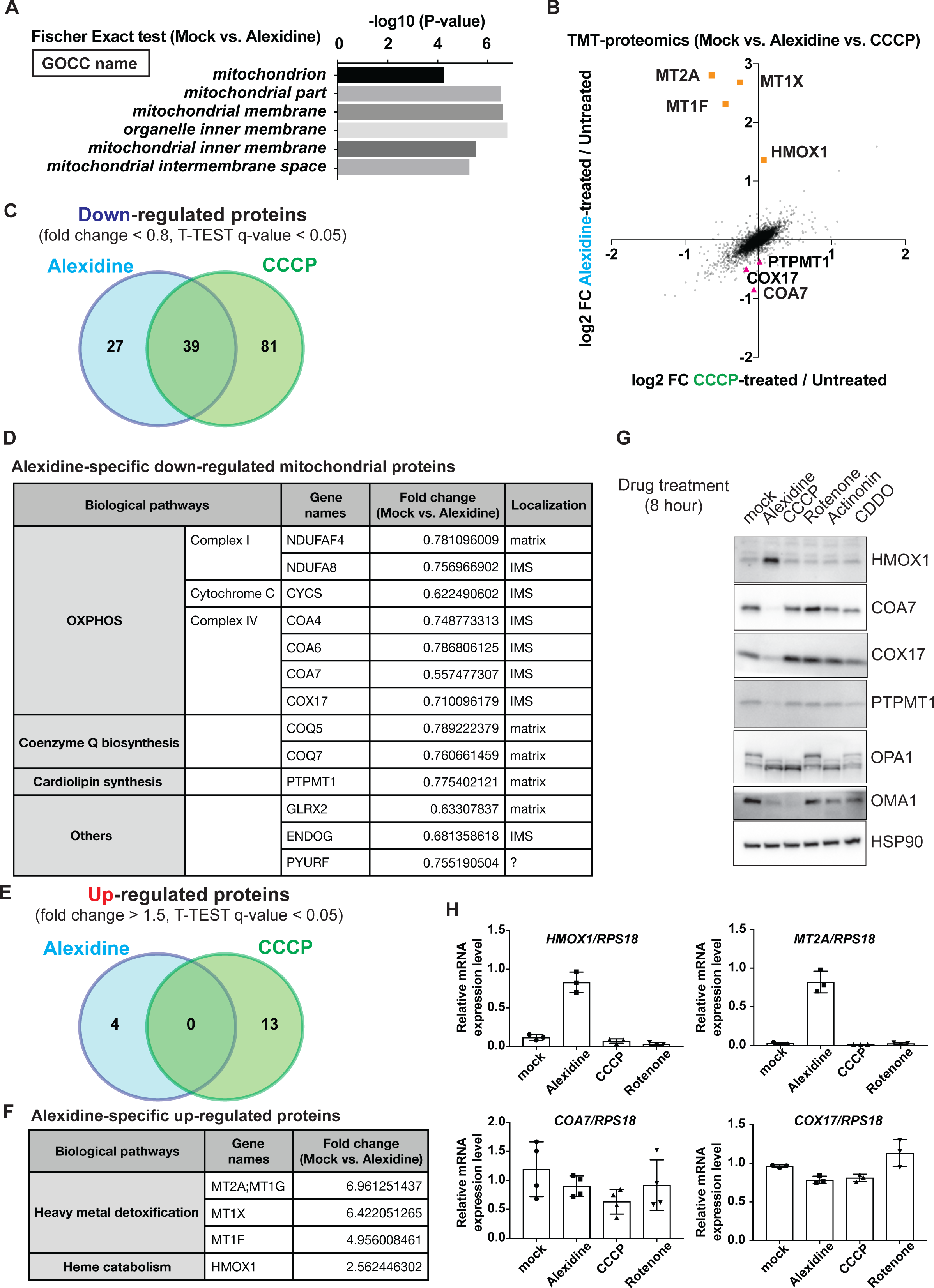
Alexidine evokes a unique transcriptional/proteostasis signature. (A-F) WT HeLa cells were treated with 5 μM Alexidine or 20 μM CCCP for 8 hours and were subjected to TMT-based quantitative proteomics. (A) The enrichment analysis of proteins whose amount was significantly changed (T-TEST q-value < 0.05) in the Alexidine-treated cells compared to the mock-treated cells. The enriched Gene Ontology Cellular Component (GOCC) classes and each enrichment value was shown. (B) Scatter plot for Log_2_ FC of protein amount in CCCP-treated/Untreated (x-axis) and Log_2_ FC of protein amount in Alexidine-treated/Untreated (y-axis). Values are from S2 table. (C) Venn diagram of down-regulated proteins (fold change < 0.8, T-TEST q-value < 0.05) after the Alexidine or CCCP-treatment. (D) List of the Alexidine-specific down-regulated mitochondrial proteins. IMS; inner membrane space. (E) Venn diagram of up-regulated proteins (fold change > 1.5, T-TEST q-value < 0.05) after the Alexidine or CCCP-treatment. (F) List of the Alexidine-specific up-regulated proteins. (G, H) Validation of TMT-based proteomics results by SDS-PAGE (G) and RT-PCR (H). WT HeLa cells were treated with the indicated drugs for 8 hours (5 μM Alexidine, 20 μM CCCP, 10 μM Rotenone, 150 μM Actinonin, and 10 μM CDDO) and were subjected to further analyses. Data are shown as mean ± S.D. (n=3 or n=4 per condition) (H).

## Discussion

In this study, we found two bactericides, Chlorhexidine and Alexidine, as small molecules that can induce the acute perturbation of mitochondrial membrane integrity. EM analysis of mitochondria after Alexidine-treatment showed a strikingly altered cristae membrane structure, while keeping the OMM and the IBM largely in place. This suggest that Alexidine can specifically damage the cristae among several distinct membrane compartments within mitochondria (Fig. 3A). It has been noted that cristae membranes are the membranes where OXPHOS proteins are concentrated [4]. Our TMT-based quantitative proteomics analysis indicated that many OXPHOS proteins were down-regulated after Alexidine-treatment (Fig. 4D), which also supports the specific action of Alexidine on the cristae membrane. Interestingly, several recent studies identified Alexidine (and Chlorhexidine) as an agent that can alter cellular metabolism. This metabolic shift ultimately resulted in various effects on cells; the anti-invasive and metastatic activity on tumor cells [60, 61], the maintenance of the quiescent status of stem cells [62], the enhanced glucose utilization *in vivo* [63], or the transcription factor TFEB nuclear translocation through the AMPK activation [64]. Some of these reports found that the Alexidine-treatment reduced oxygen consumption [62], and preferentially shifted the energy source from OXPHOS to glycolysis [61]. Because PTPMT1 was reported as a metazoan target of Alexidine [35], some studies indicated above speculated that the observed metabolic effects may result from the PTPMT1 inactivation. However, the cristae-membrane disrupting activity of Alexidine, which we identified in this study, must now be considered as a basis for the acute effects of Alexidine on the cellular metabolism.

It still remains elusive how Alexidine specifically affects the cristae membrane. As predicted from the chemical properties of guanidium groups of Alexidine [44, 45], our NAO staining assay indicated that mitochondrial membrane potential can be a driving force for the mitochondrial targeting of Alexidine (Fig. 2 D and E). Because of the high abundance of OXPHOS proteins in the cristae [4], the cristae membranes have higher membrane potential than the IBM [39]. This unique feature of the cristae may explain the specific effect of Alexidine on this membrane compartment within the IMM. Also, it has been suggested that the high-curvature of the cristae is created by high amounts of non-bilayer lipids such as CL and PE [1, 4]. We demonstrated that Alexidine has a reasonable affinity for CL as it is able to compete with NAO (Fig. 2C). Therefore, this property of Alexidine may also contribute to the accumulation of Alexidine in the cristae membrane.

As a result of Alexidine-treatment, we observed a robust induction of HMOX1, a heme-degrading enzyme, and several MTs, metal chelators (Fig. 4 F-H). The direct link between the Alexidine-mediated mitochondrial membrane damage and the induction of HMOX1 and MTs is not known. Early studies indicate that HMOX1 and MTs are simultaneously induced by heme addition to the culture media [65]. Subsequent studies have demonstrated that HMOX1 induction was mediated by nuclear factor erythroid 2-related factor 2 (NRF2), a transcription factor involved in the antioxidant response [59]. Mitochondria are known as a site for heme biosynthesis [66]. Also, the mitochondrial matrix has a pool of the heavy metal copper [66]. Together with Fe-S clusters that are synthesized in mitochondria, heme and copper are utilized as important cofactors for various enzymes including OXPHOS proteins. Due to their harmful radical-formation activity, the export of newly synthesized heme across the mitochondrial membranes is tightly regulated by a membrane-embedded heme exporter, while copper chaperones ensure the safe delivery of copper to target proteins [66]. However, it is possible that the Alexidine-mediated IMM perturbation disturbed this regulation and resulted in the heme and copper leakage from mitochondria. Ultimately, it might lead to the induction of HMOX1 and MTs as a preventive strategy. Complex IV utilizes a heme-copper center to reduce oxygen [66]. We observed that the Alexidine-treatment strongly degraded two Complex IV assembly factors; COA7 and COX17 [66–68] (Fig. 4G). The exact role of COA7 in the assembly of Complex IV is not known. Meanwhile, COX17 is well-known as a copper chaperone that delivers copper to Complex IV [66]. Therefore, in addition to the direct leakage of heme and copper from the mitochondria, it is also possible that heme and copper released from degraded OXPHOS proteins activate the transcription of HMOX1 and MTs. In either case, the induction of HMOX1 and MTs can be used as a sensitive marker of the mitochondrial membrane damage.

In addition to these transcriptional/proteostasis alterations, Alexidine remodeled the IMM-resident membrane proteins including PHB2, OMA1 and Mic60 (Fig. 3 B and D). The single particle electron microscopic analysis of PHB complex suggested that it forms a ring-like structure in the IMM [47]. It is predicted that the ring-like PHB complex can exert a partition-like function in the IMM, where it can define the lateral distribution of specific lipids, including CL and PE, or proteins such as IMM-resident proteases including OMA1 [48]. As Alexidine showed an affinity to CL (Fig. 2C), Alexidine might be able to accumulate in the PHB complex-organized CL/PE-enriched domain of the IMM. Previous reports revealed that PHB deletion can activate OMA1 proteolytic activity without any obvious mitochondrial membrane depolarization [52, 53, 69], indicating that PHB complex may hold OMA1 in the inactive state presumably through restricting the protease to specific IMM microdomains. We observed the segregation of OMA1 from PHB2-positive IMM after Alexidine treatment (Fig. 3B). These observations may indicate that Alexidine causes OMA1 to dissociate from the PHB complex-organized microdomain and that once OMA1 is released from PHB complex-mediated inhibition, it is proteolytically active. However, OMA1 did not cleave or degrade certain substrates such as PINK1 (C125G), PGAM5, CHCHD2 and CHCHD10 upon Alexidine treatment (Fig. 1K). This observation implies that the PHB complex-organized lipid microdomain not only regulates the proteolytic activity of OMA1 but may also be involved in the spatial regulation of OMA1 and its substrates.

Mic60, one of the important components of MICOS complex [46], showed a puncta-like localization within the IMM after Alexidine treatment (Fig. 3D). Among several components of MICOS complex, it has been suggested that Mic60 can self-assemble and forms puncta within the IMM when all other MICOS components are absent in yeast [56]. Moreover, the recent analysis showed the *de novo* formation of CJs by drug-controlled expression of Mic60 in reconstituted Mic60 KO cells [70], establishing a critical role for Mic60 in CJ formation. Upstream determinants of Mic60 localization are not fully understood. However, recent studies in yeast has revealed that the aforementioned Mic60 puncta formed in the absence of other components of the MICOS complex are often observed in proximity to ER-mitochondria contact sites, where the ERMES complex exists [71]. The ERMES complex physically tethers the ER and mitochondria in yeast and creates membrane contact sites to allow efficient lipid transfer between the ER and mitochondria [2]. Together with the fact that MICOS and ERMES genetically interact with each other [55], it has been suggested that cooperative functions of MICOS and ERMES in mitochondrial membrane architecture (ERMIONE) [72]. Mitochondrial lipid homeostasis depends on both inter-organelle (mainly from the ER) and intra-organelle (between the OMM and the IMM) lipid trafficking [2, 3]. Therefore, the coordinated regulation of ERMES-MICOS localization may be functionally linked to allow for efficient lipid trafficking across the membranes. Although it has not been examined whether the MICOS complex is involved in inter-organelle lipid trafficking, it is known that the MICOS complex is involved in intra-mitochondrial lipid metabolism [57, 73]. Our EM analysis revealed that the cristae structure was severely damaged following Alexidine treatment (Fig. 3A). In such a case, it is expected that mitochondrial lipid demand is significantly increased in order to help restore the highly-folded cristae structure. It is tempting to speculate that the Alexidine-induced Mic60 puncta formation plays a role in this process as a part of a mitochondrial membrane stress response. It may be also of note that several reports indicate a link between the MICOS complex and COA7 or COX17 (robustly degraded proteins after the Alexidine-treatment). For example, in yeast, it was shown that COX17 physically interacts with Mic60 and modulates the MICOS complex formation [74]. Other studies suggest that the sustained KD of Mic60 (or its interactor, Sam50) resulted in the degradation of COA7 in mammalian cells [68]. COA7 was also identified as the possible interactor of Mic10, another important component of MICOS complex [75]. These previous observations may explain the reason why we observed the relatively specific and robust degradation of COA7 and COX17 among over 100 components of OXPHOS system.

In conclusion, we discovered Alexidine and Chlorhexidine as small molecules that enable us to acutely and preferentially perturb the mitochondrial membrane architecture in the IMM. Our findings therefore offer a useful chemical-biological tool for delineating mitochondrial membrane stress responses.

---

**Supplementary Figure 1.**
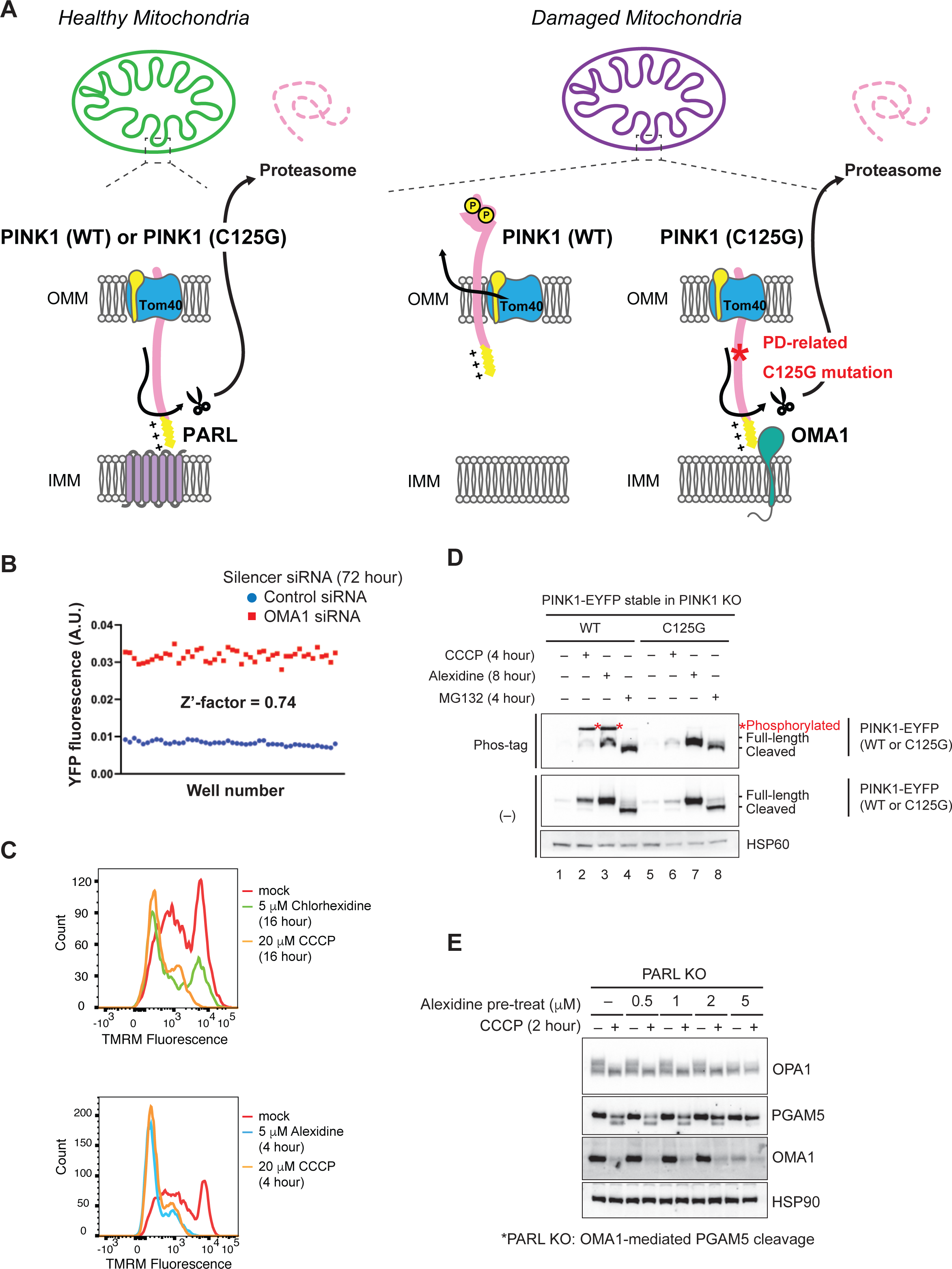
Compound screen of OMA1 inhibitors identifies bactericides. (A) The mitochondrial import pathway of PINK1. See text for detail. (B) PINK1 KO HeLa cells stably expressed PINK1 (C125G)-EYFP were seeded on 96 well plate and were transfected with control or OMA1 siRNA (48 wells for each). After 72 hours, cells were treated with 20 μM CCCP for 4 hours. Z’-factor (control siRNA vs. OMA1 siRNA) was calculated by using EYFP fluorescence intensity measured by a high content image analyzer. (C) WT HeLa cells were treated with the indicated drugs for the indicate time period. After each drug-treatment, mitochondrial membrane potential of each cell was measured by FACS using tetramethylrhodamine-methyl ester (TMRM). (D) PINK1 KO HeLa cells stably expressed with PINK1 wild-type (WT)-EYFP or PINK1 (C125G)-EYFP were treated with the indicated drugs for the indicated time period. The lysate was analyzed by SDS-PAGE with or without Phos-tag. (E) PARL KO HeLa cells were pre-treated with Alexidine at the indicated concentration for 30 min, and after, treated with 20 μM CCCP for 2 hours. Note that in PARL KO cells, the CCCP-dependent PGAM5 cleavage is mediated by OMA1. The lysate was analyzed by SDS-PAGE.

**Supplementary Figure 2.**
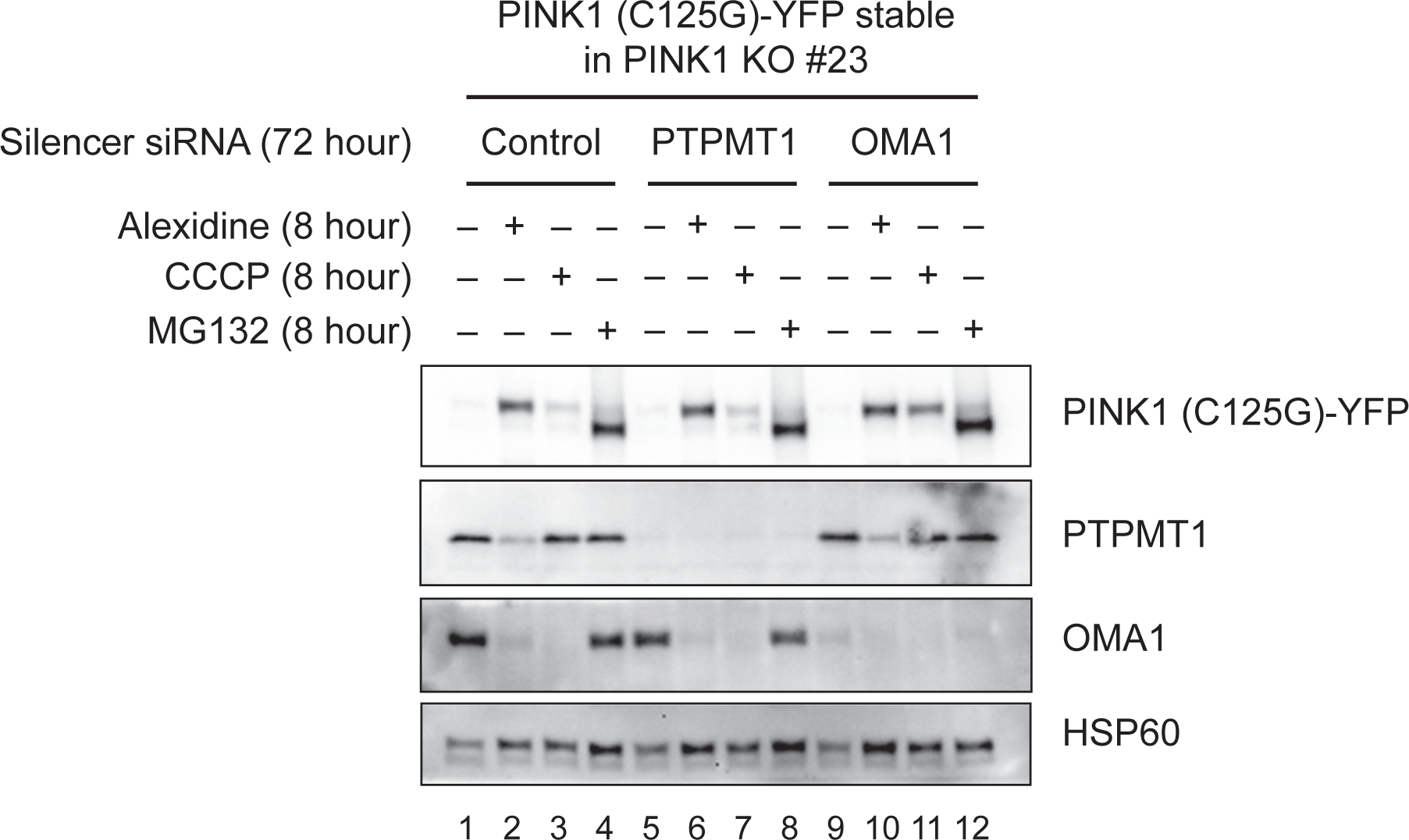
Transient deletion of PTPMT1 did not promote the accumulation of PINK1 (C125G). PINK1 KO HeLa cells stably expressed PINK1 (C125G)-EYFP were transfected with control, PTPMT1 or OMA1 siRNA. After 72 hours, cells were treated with the indicated drugs for 8 hours. The lysate was analyzed by SDS-PAGE.

**Supplementary Figure 3.**
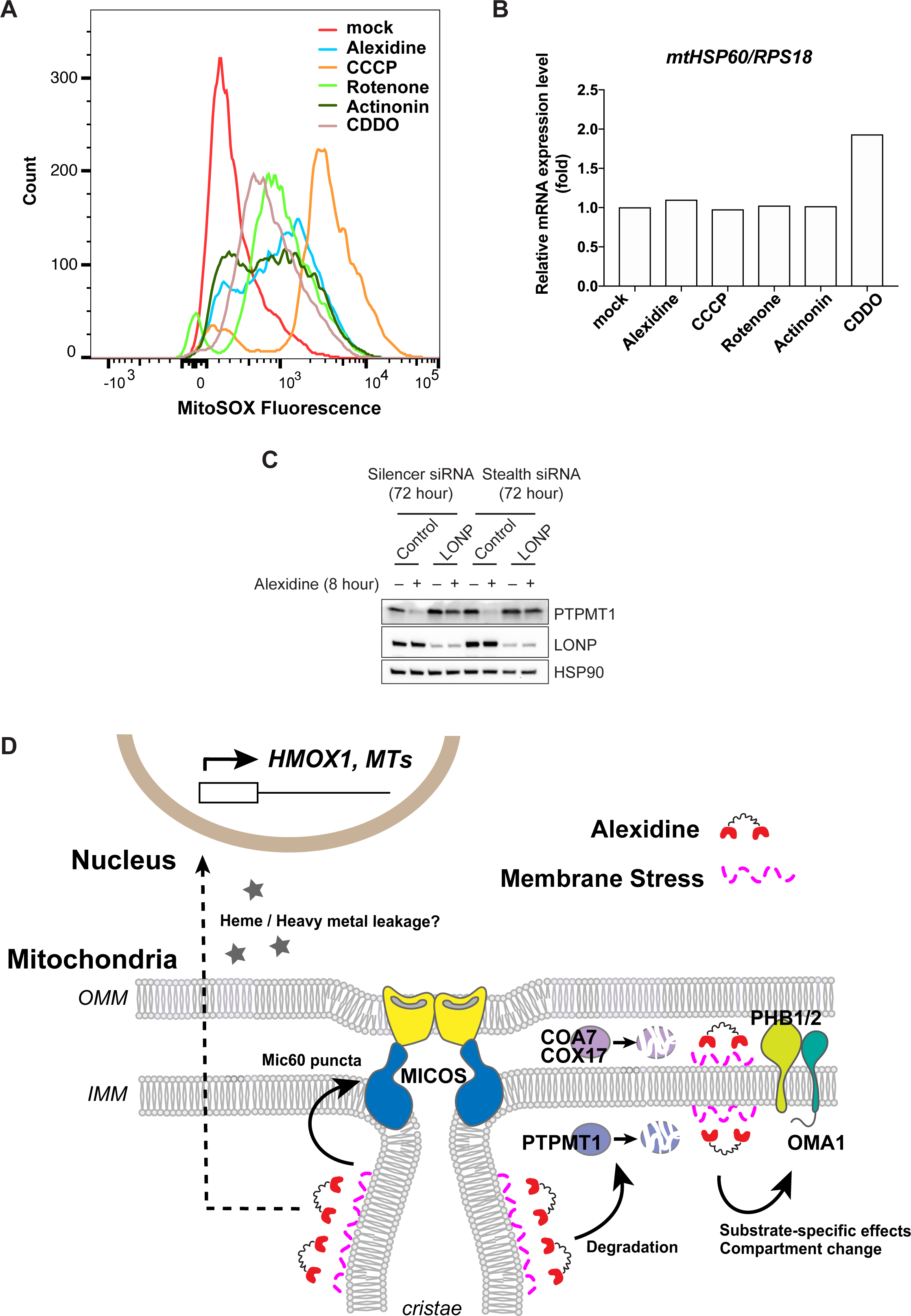
Alexidine is a novel mitochondrial stressor that can induce the acute mitochondrial membrane damage. (A) WT HeLa cells were treated with the indicated drugs for 4 hours. After each drug-treatment, mitochondrial ROS production of each cell was measured by FACS using MitoSOX. (B) WT HeLa cells were treated with the indicated drugs for 4 hours (5 μM Alexidine, 20 μM CCCP, 10 μM Rotenone, 150 μM Actinonin, and 10 μM CDDO) and were subjected to RT-PCR analysis. (C) WT HeLa cells were transfected with the indicated control or LONP siRNAs. After 72 hours, cells were treated with 5 μM Alexidine for 8 hours and were subjected to SDS-PAGE. (D) Graphical summary of this study. See text for detail.

**Supplementary Table 1 The results of the FDA-approved compound screen.**

**Supplementary Table 2 The results of the TMT-based quantitative proteomic analysis.**

## Materials and methods

### Cell culture, transfection, and treatments

HEK293T and HeLa cells were cultured in DMEM (Gibco) supplemented with 10% FBS (VWR Life Science), 10 mM HEPES (Gibco), 1 mM Sodium pyruvate (Gibco), non-essential amino acids (Gibco) and GlutaMAX (Gibco). Following HeLa KO cell lines were kindly provided by Dr. Richard J Youle (NIH, NINDS); PINK1 KO [76], OMA1 KO [23], and PARL KO [23]. For RNA interference, 20 nM Stealth siRNAs (Steath RNAi Negative Control Med GC Duplex #2, 12935112; LONP, HSS113887) (Thermo Fisher Scientific) or 5 nM Silencer select siRNAs (Silencer Select Negative Control #1 siRNA, 4390843; OMA1, s41776; LONP, s17903; PTPMT1, s229947) (Thermo Fisher Scientific) were transfected using Lipofectamine RNAi max transfection reagent (Thermo Fisher Scientific) at the same time as cell seeding. For drug treatment experiments, cells were incubated in medium containing one or more of the following compounds: CCCP (Cayman Chemical), MG132 (Sigma), Chlorhexidine (Cayman Chemical), Alexidine (Cayman Chemical), Rotenone (Cayman Chemical), Actinonin (Cayman Chemical), CDDO (Cayman Chemical). For examination of mitochondrial membrane potential or mitochondrial ROS production, 20 nM TMRM (Thermo Fisher Scientific) or 50 nM MitoSOX (Thermo Fisher Scientific) respectively was directly added to cell culture media and incubated for 15 min. For NAO staining, cells were washed twice with PBS and were incubated with 100 nM NAO (Thermo Fisher Scientific) for 15 min. Cells were washed and replaced with normal medium followed by live cell imaging using a 63 × /1.4 NA oil immersion objective on Leica SP8 LIGHTNING Confocal Microscope (Leica) or FACS analysis using Attune NxT Acoustic Focusing Cytometer (Thermo Fisher Scientific).

### Plasmids

Site-directed mutagenesis of pLenti-CMV-Neo-PINK1 (C125G)-EYFP or pLVX-puro-OMA1 (E328Q)-EYFP was performed by PCR amplification (CloneAmp HiFi PCR Premix, Takara or Q5 High-Fidelity DNA Polymerase system, NEB) of PINK1 or OMA1 encoding plasmid using appropriate primers followed by Gibson assembly (In-Fusion HD Cloning system, Clontech) into the SalI-XhoI of the pLenti-CMV-Neo vector, or into the EcoRI site of pLVX-puro vector (Clontech). pLVX-puro-DELE1-HA and pLVX-puro-Su9-mCherry was created by PCR amplification and sub-cloning into the EcoRI site of pLVX-puro vector. All constructs were confirmed by DNA sequencing.

### Generation of stable cell lines

To generate stably transfected cell lines, lentiviruses (for plasmids within pLenti-CMV-neo or pLVX-puro vectors) were packaged in HEK293T cells. HeLa cells were transduced with viruses with 10 μg/ml polybrene (Sigma) then optimized for protein expression via antibiotics selection. PINK1 KO HeLa cells stably expressed PINK1 (C125G)-EYFP were mono-cloned, and clone #23 was used in this study.

### Immunoblotting (IB)

For SDS-PAGE, cells were lysed with 1× NuPAGE LDS sample Buffer (Thermo Fisher Scientific) supplemented with 100 mM Dithiothreitol (DTT) (Sigma), and boiled with shaking for 15 min. Approximately, 50 μg of protein per sample was separated on 7.5 %, 10 % or 4-20% gradient Mini-PROTEAN TGX Precast Gel (Bio-Rad) or Criterion TGX Gels (Bio-Rad) and then transferred to a nitrocellulose membrane (Bio-Rad) or PVDF membrane (Bio-Rad). The membrane was blocked with Odyssey Blocking Buffer (LI-COR) and incubated with the indicated primary antibodies at 4°C overnight. After washing with PBS-T (PBS + 0.05% Tween-20), the membrane was incubated with HRP-conjugated secondary antibodies (Thermo Fisher Scientific) and washed again with PBS-T. Detection was performed with iBright CL1000 Imaging System (Thermo Fisher Scientific). For Phos-tag SDS-PAGE, cells were lysed with 1% Triton buffer [1% Triton-X100, 150 mM NaCl, 50 mM Tris-HCl pH 7.4, 1 mM EDTA, Phosphatase inhibitors (PhosSTOP, Sigma) and protease inhibitors (cOmplete, Sigma)]. After centrifugation, the lysate that contains 10 μg of protein per sample was mixed with 2x Laemmli Sample Buffer (Bio-Rad) supplemented with 2 M 2-Mercaptoethanol (Bio-Rad), and boiled for 3 min. Samples were separated on 7.5 % Mini-Gel (TGX FastCast Acrylamide Solutions, Bio-Rad) containing 50 μM Phos-tag AAL-107 (Wako) and 10 mM MnCl_2_ (Sigma) according to the manufacturer’s instruction. For the elimination of the manganese ion from the gel, the gel was soaked with a transfer buffer containing 1 mM EDTA for 10 min, and washed with a transfer buffer without EDTA for 10 min, and then transferred to a PVDF membrane (Bio-Rad).

### Immunofluorescent chemistry (IF)

Cells were seeded into Lab-Tek Chambered Coverglass with 4 wells (Thermo Fisher Scientific). Cells were rinsed in PBS and fixed with PBS containing 4% formaldehyde for 15 min at room temperature. Cells were permeabilized with PBS containing 0.1% Triton X-100 for 10 min at room temperature. Blocking was performed using PBS containing 2% BSA for 30-60 min at room temperature. For immunostaining, cells were incubated with primary or secondary antibodies (Alexa Fluor, Thermo Fisher Scientific) diluted in PBS containing 2% BSA for overnight at 4C° or about 2 hours at room temperature, respectively. Cells were imaged using a 63 × /1.4 NA oil immersion objective on Leica SP8 LIGHTNING Confocal Microscope (Leica).

### Antibodies

Following antibodies were used in IB or IF; PINK1 (Cell Signaling, 6946S), OMA1 (Santa Cruz, sc-515788), PGAM5 (Thermo Fisher Scientific, PA5-57894), CHCHD2 (Proteintech, 66302-1-lg), CHCHD10 (Sigma, HPA003440), OPA1 (BD Biosciences, 612607), HSP90 α/β (Santa Cruz, sc-7947 or sc-13119), HSP60 (Santa Cruz, sc-13115), Tim50 (Santa Cruz, sc-393678), Tom20 (Santa Cruz, sc-17764), PHB2 (Proteintech, 66424-1-lg), Cox IV (Thermo Fisher Scientific, PA5-19471), Mic60 (Proteintech, 10179-1-AP), HMOX1 (GeneTex, GTX101147), PTPMT1 (Santa Cruz, sc-390901), COX17 (Proteintech, 11464-1-AP), COA7 (Proteintech, 25361-1-AP), LONP (Novus Biologicals, NBP1-81734), GFP (Novus Biologicals, NB600-597 or NB-600-308), and HA (Cell Signaling, 3724).

### FDA-Approved Compound Library Screening

The FDA-approved compound library (Selleck, 100 nL per drug) was stamped to black 384 well plates with glass bottom using CyBio Well vario (Analytik Jena). PINK1 KO HeLa cells stably expressed PINK1 (C125G)-EYFP (clone #23) were then added to give density of 4000 cells per well and a final drug concentration of 5 µM. After 18 hours of treatment, CCCP was added to a final concentration of 20 µM and incubated for 4 hours followed by fixation in 4% paraformaldehyde and counterstaining with Hoechst 33342. Fluorescence was detected using an ImageXpress Micro XLS (Molecular devices) high-content imager and cellular EYFP fluorescence signal was calculated using CellProfiler software [77].

### In vitro NAO assay

Binding of NAO to each anionic phospholipid was studied in lipid monolayers [42, 43], with slight modifications. 96 well microtiter plates (Corning, 3915) were coated with 50 µL of 20 µM each anionic phospholipid in ethanol and evaporated at 37°C for 5 hours. Increasing concentration of Alexidine (final; 0-100 µM) or CCCP (final; 10 µM) in 50 µL PBS containing 2% BSA was added and incubated at 37°C for 30 min. Then, NAO (final; 100 nM) in 50 µL PBS containing 2% BSA was added and incubated at 37°C for 30 min protected from light. After the incubation, each well of the plates was washed with 150 µL PBS for 5 times. Finally, NAO fluorescence intensity was measured by SpectraMax i3x Multi-Mode Detection Platform (Molecular Devices) using Ex 485nm /Em 535 nm. Following anionic phospholipids purchased from Avanti Polar Lipids were used; Heart CA (840012P), Egg PA (840101P), Egg PG (841138P), Liver PI (840042P), and Brain PS (840032P).

### Transmission Electron Microscopy

HeLa cells were fixed with 4% glutaraldehyde (Electron Microscopy Services) in EM buffer (0.1 N sodium cacodylate at pH 7.4 with 2 mM calcium chloride) for 30 min at room temperature and then at 4°C for at least 24 hours. Samples were washed with buffer and treated with 1% osmium tetroxide in 0.1 N cacodylate buffer at pH 7.4 for 1 hour on ice, washed and *en bloc* stained with 0.25–1% uranyl acetate in 0.1 N acetate buffer at pH 5.0 overnight at 4°C, dehydrated with a series of graded ethanol and finally embedded in epoxy resins. Ultrathin sections (70 nm) were stained with lead citrate and imaged with a JEOL 1200 EXII Transmission Electron Microscope.

### RNA Isolation and Real-Time PCR

Total RNAs were isolated using RNeasy Mini Kit (Qiagen) and reverse-transcribed to cDNA using iScript cDNA Synthesis Kit (Bio-Rad) according to the manufacturer’s instruction. Real-Time (RT) PCR was performed using SYBR Green Master Mix (Bio-Rad) and QuantStudio 3 RT-PCR system (Thermo Fisher Scientific). All expression levels were normalized to that of RPS18 mRNA. The following RT-PCR primers were used; RPS18, (forward) 5’- cttccacaggaggcctacac-3’ and (reverse) 5’- cgcaaaatatgctggaacttt-3’; HMXO1, (forward) 5’- ggcagagggtgatagaagagg-3’ and (reverse) 5’- agctcctgcaactcctcaaa-3’; MT2A, (forward) 5’- aacctgtcccgactctagcc-3’ and (reverse) 5’- Gcaggtgcaggagtcacc-3’; COA7, (forward) 5’-gcaggtcaagtcctttttgg-3’ and (reverse) 5’- ccaccagccgatagcaac-3’; COX17, (forward) 5’-aagatgccgggtctggtt-3’ and (reverse) 5’- ttcttctcctttctcgatgataca-3’; mtHSP60, (forward) 5’-cctgcactctgtccctcact-3’ and (reverse) 5’- gggtaaccgaagcatttctg-3’.

### TMT-based quantitative proteomics

*Sample Preparation* – Cells were lysed in 8 M urea, 50 mM Tris-HCl (pH 8.0), 1X Complete Protease Inhibitor (Roche) and 1X PhosStop (Roche) with a sonic probe (3 x 30s at 80% amplitude) with subsequent mixing at room temperature for 1 hour at 1000 rpm on a Thermomixer. Lysates were quantified by Qubit fluorometry (Life Technologies). 50 µg of each sample was digested overnight with trypsin. Briefly, samples were reduced for 1 hour at room temperature in 12 mM DTT followed by alkylation for 1 hour at room temperature in 15 mM iodoacetamide. Trypsin was added to an enzyme:substrate ratio of 1:20. Each sample was acidified in formic acid and subjected to SPE on an Empore SD C18 plate. Each sample was lyophilized and reconstituted in 140 mM HEPES (pH 8.0), 30% acetonitrile for TMT labeling. Peptides were labeled using TMT 10-plex (Thermo Fisher Scientific) according to manufacturer’s instructions. Briefly, 40 µL of acetonitrile was added to each TMT tag tube and mixed aggressively. Tags were incubated at room temperature for 15 min. 30 µL of label was added to each peptide sample and mixed aggressively. Samples were incubated in an Eppendorf Thermomixer at 300 rpm at 25°C for 1.5 hour. Reactions were terminated with the addition of 8 µL of fresh 5% hydroxylamine solution and 15 min incubation at room temperature. Each labeled sample was pooled, frozen, and lyophilized and subjected to SPE on a High-Density 3M Empore SDB-XC column. The eluent was lyophilized. Peptides were fractionated using high pH reverse-phase chromatography on an Agilent 1100 HPLC system using a Waters XBridge C18 column (2.1mm ID x 150mm length, 3.5µm particle size) at 300 µL/min. The following gradient was employed: 0.5% B initial conditions, 0.5-3.0% B from 0-1 min, 3-25% B from 1-36 min, 25%–45% B from 36-44 min, 45-90% B from 44-47 min, 90% B from 47-49 min, 90%–0.5% buffer B from 49-50 min (buffer A: 100% H_2_O, 10 mM NH_4_OH; buffer B: 100% CH_3_CN, 10 mM NH_4_OH). Every 12^th^ well was combined to create 12 pools. Each pool was lyophilized. *Mass Spectrometry* - Peptides (10% per pool) were analyzed by nano LC/MS/MS with a Waters NanoAcquity HPLC system interfaced to a ThermoFisher Fusion Lumos mass spectrometer. Peptides were loaded on a trapping column and eluted over a 75 µm analytical column at 350 nL/min; both columns were packed with Luna C18 resin (Phenomenex). Each high pH RP pool was separated over a 2 hours gradient (24 hours instrument time total). The mass spectrometer was operated in data-dependent mode, with MS and MS/MS performed in the Orbitrap at 60,000 FWHM resolution and 50,000 FWHM resolution, respectively. A 3 s cycle time was employed for all steps. *Data Analysis* – Data were analyzed using MaxQuant v1.6.2.3 (Max Planck) and searched against the combined forward and reverse Swissprot *H. sapiens* protein database. The database was appended with common background proteins. Search parameters were precursor mass tolerance 7 ppm, product ion mass tolerance 20 ppm, 2 missed cleavages allowed, fully tryptic peptides only, fixed modification of carbamidomethyl cysteine, variable modifications of oxidized methionine and protein N-terminal acetylation. Data were filtered 1% protein and peptide level false discovery rate (FDR) and requiring at least one unique peptide per protein. Reporter ion intensities were exported for further analysis.

### Statistical analysis

Statistical significances were determined using Prism software (GraphPad Software, Inc.) as indicated in the Figure legends.

## Acknowledgements

We thank Dr. Toren Finkel for critical reading our manuscript and thank Dr. Richard J Youle for sharing materials. We thank Dr. Susan Cheng, Ms. Sandra Lara, and the NINDS, EM Facility (NIH) for technical assistance with transmission electron microscopy. We thank Dr. Richard Jones and MS Bioworks, LLC for performing TMT-based quantitative proteomics. This work was supported by University of Pittsburgh, Aging Institute Startup Seed, Samuel and Emma Winters Foundation, and UPMC Health System Competitive Medical Research Fund. This work was supported (in part) by the NINDS intramural program.

## Author contributions

R.H., Y.S., and S.S. designed and performed experiments and wrote the manuscript. M.L. and B.B.C. performed the FDA-approved compound screening. K.M. provided chemical-structural insights on hit compounds. D.P.N. designed the EM analysis and interpreted the EM images.

## Conflict of interest

The authors declare no competing financial interest.

## Notes

### Competing Interest Statement

The authors have declared no competing interest.

